# Evaluation of selectively-activatable, caged fluorescent probes as species selective markers for beta-alanine aminopeptidase positive bacterial species

**DOI:** 10.64898/2026.06.28.734737

**Authors:** Louise Soh, Mohammad Askarzadeh, Khondaker Miraz Rahman, Charlotte K. Hind, J. Mark Sutton

## Abstract

Aminopeptidases are widely distributed in bacteria, but outside of a few model strains, their function is largely unexplored. Focussing on beta-alanine aminopeptidase activity, a new series of selectively-activatable, caged fluorescent probes were designed and synthesised. A beta alanine amino acid was coupled to resorufin or 7’-hydroxycoumarin via a self-imolative linker, such that amino acid removal led to gain of fluorescence. These were used to probe selectivity and specificity of probe activation, against a range of priority drug-resistant pathogens. When added to bacterial growth curves run in Muller Hinton broth, these probes allowed essentially real time fluorescence measurement of activation by bacterial species, modelled on the standard microbroth dilution method. Activation was observed for all *Pseudomonas aeruginosa* and Burkholderia spp strains tested. Selective activation was seen for Ochrabactrum species, with the probe activated by *O.anthropii* (2/4 strains) but not *O.intermedium* and strain-specific activation was seen for some isolates of *Serratia marcescens* (2/4 strains). No activation was observed in any isolates of *Klebsiella pneumoniae, Escherichia coli, Acinetobacter baumannii* or *Staphylococcus aureus* or *Eneterocccus faecium/faecalis*

PAO1 transposon mutants in the putative beta-alanine aminopeptidase gene (annotated as *bapF* or *dmpA*; PW3678) showed no activation of the probe in growth assays, confirming the specificity of the probe for beta-alanine aminopeptidase. Transposon mutants in other aminopeptidase genes, including those encoded by *pepN, PepP* and the prolyl aminopeptidase gene had no effect on probe activation in PAO1. Based on the operon structure in PA01, transposon mutants in two adjacent genes were also tested for probe activation. Mutants in both a putative transcriptional regulator (PW3674) and a predicted amino acid permease (PW3676) retained their ability to activate the beta-alanine probes with activation significantly higher than the wild type, when assessed by the total fluorescence yield after 10 hours growth. This points to both redundancy in permease function and perhaps the presence of a feedback regulatory mechanism controlling beta alanine aminopeptidase activity in *P.aeruginosa*. Given that the operon structure is conserved in other species, this may point to a common mechanism of beta alanine aminopeptidase function, perhaps related to exploiting beta-alanine containing peptides in certain environmental niches.

## Introduction

### 1.2 Bacterial enzymes as biomarkers

There is a significant unmet need for diagnostic tests and biomarkers that are rapid, reliable, and easy to use and can be applied in the detection and diagnosis of infectious disease. Rapid diagnostic technologies can play a crucial role in improving the accuracy of bacterial identification and supporting the appropriate prescription of antibiotics. Their implementation could facilitate a shift towards the use of targeted narrow-spectrum antibiotics rather than broad-spectrum agents, thereby helping to reduce the emergence and spread of antimicrobial resistance. Species-specific biomarkers can be used to understand the importance of specific bacteria in mixed communities.

Despite these advantages, many current diagnostic methods are still performed in clinical laboratories using phenotypic, culture-based, or genotypic approaches. These methods are often limited by lengthy turnaround times or the requirement for sophisticated laboratory infrastructure, expensive instrumentation, and highly trained personnel. Phenotypic diagnostic methods frequently rely on biochemical assays that assess enzymatic activity or metabolic characteristics for bacterial identification. Considerable research interest has focused on the detection of metabolic substrates as indicators of enzymatic activity, providing valuable tools for the identification and characterisation of bacteria. Typically, synthetic enzymatic substrate complexes consist of a target substrate linked to a reporter molecule, such as a dye or fluorophore. Following an enzymatic reaction, commonly hydrolysis, the reporter is released, generating a measurable signal. The reporter is essential because it enables enzymatic activity to be detected through a visible or quantifiable response. Chromogenic, fluorogenic, and luminogenic substrates are widely used for this purpose. In some instances, enzymatic activity can also be detected directly through the measurement of naturally produced metabolic products.

A wide range of bacterial enzymes have been investigated and utilised as taxonomic markers for microbial identification. Enzyme classes including esterases, glycosidases, arylamidases, nitroreductases, and aminopeptidases have been successfully employed for the detection, identification, and enumeration of bacterial species.^1^ This study focusses on bacterial aminopeptidases given the diversity in bacteria and the relatively limited study of their function in many species.

Bacterial aminopeptidases, which remove the amino terminal amino acid from a protein or peptide, can also be classified according to their substrate specificity (broad or narrow), and by the requirements they may have for specific amino acids on either side of the substrate cleavage site. Aminopeptidases that exhibit broad substrate specificity are usually incapable of hydrolysing peptide bonds formed by acidic amino acid residues (Asp, Glu) in the P1 (N-terminal amino acid) position, or proline at either P1 or P1’ (first amino acid after the cleavage site) position.^2^ While exhibiting a broad substrate specificity, some aminopeptidases has shown preference for certain amino acids. For example, *E.coli* PepN is a broad specific aminopeptidase that cleaves alanine, arginine, lysine, serine, leucine, methionine etc., with maximal activity with alanine.^3^ In colorimetric assays, *E.coli* PepA prefers substrates leucine, methionine, and also phenylalanine, arginine and alanine.^4^ Pseudomonas putida PepA is most active on leucine, methionine and also isoleucine, valine, phenylalanine.^5^ Aminopeptidases with narrow specificities may only accommodates a single type of amino acids, or less efficiently a small group of related amino acids. The prolyl aminopeptidase (PAP) is an example of narrow substrate specificity where it almost exclusively releases a proline from the N-terminus of a peptide.^6^ The activity of aminopeptidase with narrow specificity is not only determined by the amino acid residue at the N-terminus, but may also be determined by the P1’ amino acid or other primary sequence or structural features.

Aminopeptidases play a number of often overlapping roles in bacteria. For example, they may be involved in scavenging amino acids from exogenous peptides and to support uptake of nutrients from the environment. The aminopeptidase activity in the lactic acid bacteria, *Lactobacillus lactis* during the fermentation of milk products has been well studied.^7–8^ where they hydrolyse milk proteins, caseins, into amino acids suitable for nutrient uptake. Aminopeptidases found in *L. lactis* displayed a preference in cleaving proline substrates, which is adapted to its environment as caseins are rich in proline. Aminopeptidases are also involved in turnover of protein and peptides, to maintain the balance between the synthesis of new proteins and the catabolism of existing and sometimes damaged proteins. This may be linked to growth cycle, where proteins are degraded more rapidly during stationary phase than logarithmic growth phase; at least partially addressing the nutrient demands of the dividing cell. Protein turnover is also essential to eliminate abnormal or damaged proteins and cleaved signal sequence peptides that are derived from exported proteins. ^9, 2^ Aminopeptidases are responsible for catalysing the final stages of proteolysis during the degradation of endogenous proteins, following cleavage by one of more of the ATP-dependent proteases, such as ClpXP. In *Salmonella typhimurium* aminopeptidases A, B, D and N are involved in the final stages of degradation of intracellular and abnormal proteins into amino acids.^10,11^ Bacterial aminopeptidases are thought to be essential for post-translational modification and maturation of proteins. Methionine aminopeptidase MetAP, cleaves N-terminal methionine residue of newly synthesized polypeptide chains and is essential as deletion of the metAP gene(s) result in a lethal phenotype in *E.coli* and *S.typhimurium.*^12,13^ In the absence of methionine aminopeptidase, the main cell proteins remained inactivated as the methionyl precursors. Other aminopeptidases and dipeptidyl aminopeptidases are involved in interactions with the host, for example, turning over host defense peptides^14^

We focussed on the development of selectively activatable, fluorescent probes for beta-alanine aminopeptidases, a class of aminopeptidases which are relatively restricted in bacteria. The aim was to generate a novel probe that could be used for real time detection and growth monitoring in liquid (rather than in agar ^15^). This is applied to the important, WHO Priority pathogen *P.aeruginosa* to explore its specificity and selectivity for this over other pathogens. Real-time monitoring of aminopeptidase activity in intact bacterial cells using fluorescent probes provides a powerful approach for investigating the biological roles of these enzymes. Such probes enable the direct measurement of aminopeptidase activity in living bacteria, allowing the distribution, abundance, and temporal expression of aminopeptidases to be characterised across different bacterial species. Understanding these patterns is essential for assessing the potential of bacterial aminopeptidases as diagnostic biomarkers and for exploring their application in areas such as bacterial species differentiation and the development of species-specific prodrugs.^16^

The initial phase of the project centred on the design and synthesis of fluorescent probes for the direct detection of aminopeptidase activity in whole bacterial cells. A key objective was to develop highly sensitive probes capable of rapidly identifying even low levels of enzymatic activity and/or low cell numbers. To achieve this, the probes were engineered to function through a fluorescence “off-to-on” mechanism, in which enzymatic cleavage triggers a marked increase in fluorescence intensity. Compared with constitutively fluorescent systems, this approach offers significant advantages because the intact probe produces very little background fluorescence. As a result, signal generation occurs only upon enzyme-mediated activation, leading to greater sensitivity and improved signal-to-noise performance.

Figure 1 presents the general architecture of the aminopeptidase-responsive fluorescent probe as well as the mechanism of cleavage. The probe consists of three principal components: an amino acid recognition unit, a fluorophore reporter, and a *para*-aminobenzyl alcohol (PABA) self-immolative linker. The amino acid substrate acts as the recognition element for the target aminopeptidase, while the fluorophore serves as the signal-generating component. The PABA self-immolative linker is incorporated to spatially separate the amino acid substrate from the relatively bulky fluorophore, thereby reducing steric hindrance and promoting efficient interaction of the substrate with the enzyme active site. This design enhances enzymatic accessibility and facilitates effective probe activation upon aminopeptidase-mediated cleavage ^17^.

**Figure 1.**
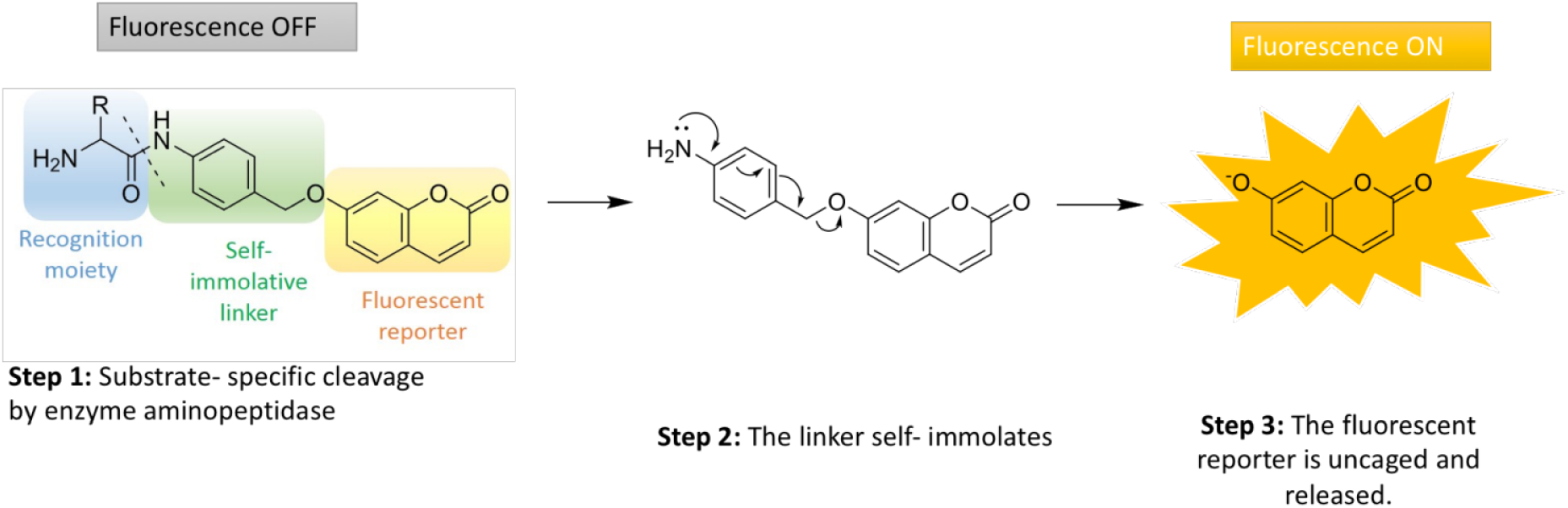
The design and mechanism of activation of an off-on fluorescent probe to detect aminopeptidase activity in bacteria.

**Figure 2.**
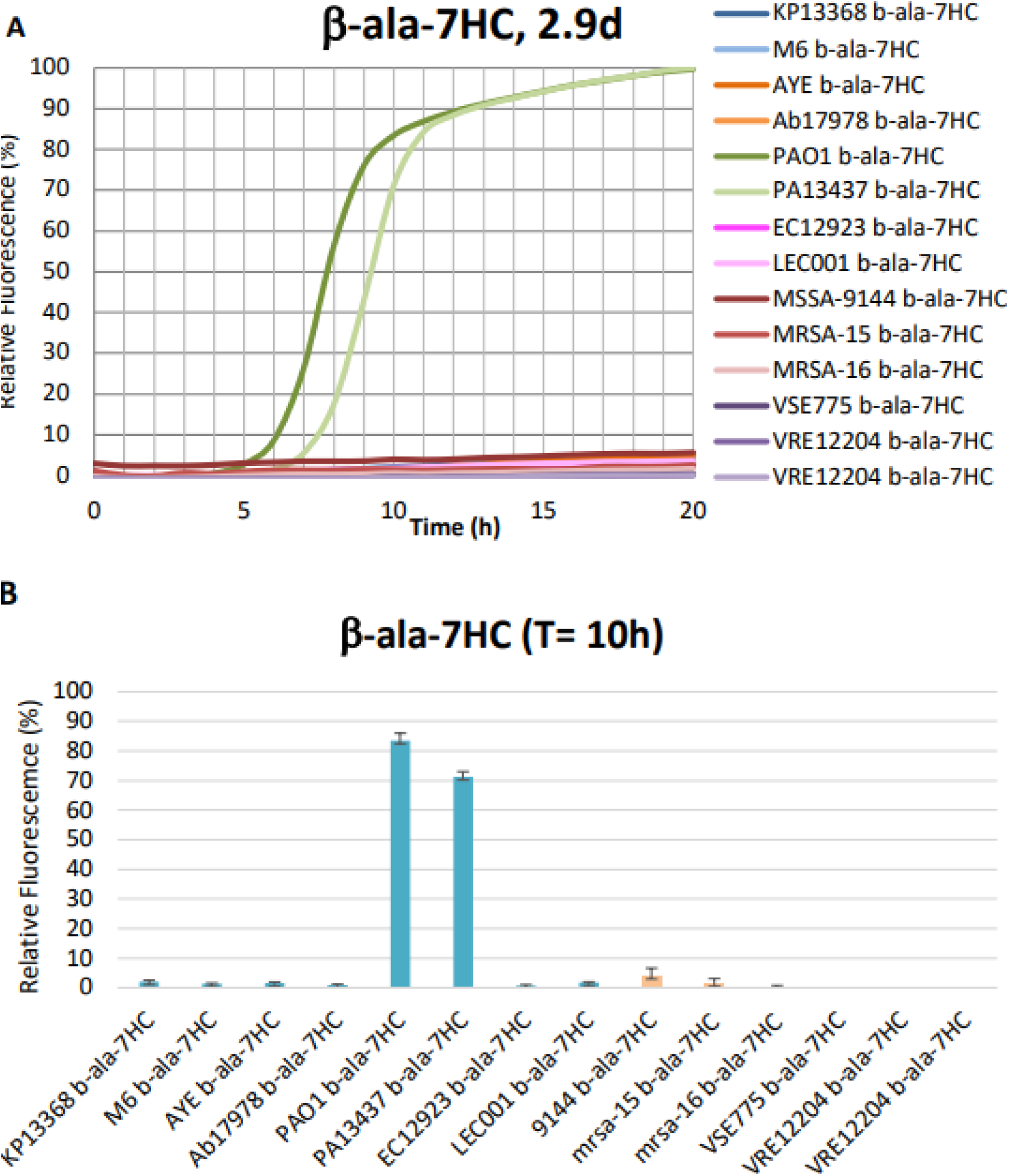
Comparison of the fluorescent probe activation assessed using a beta-alanine-7HC fluorescent probe. (A) presented as relative fluorescence (%) over time (h) graphs. The graph is a representative of at least 3 biological repeats. The fluorescence assay is performed by adding diluted overnight bacteria culture of OD600 0.01 (approximately 5x10^5^ cfu/ml) with 25 µM of each probe on a microplate. The fluorescence generated was measured at 460 nm, hourly over 20 hours at 37°C. (B) The average fluorescence activity of activated beta-alanine-7HC at 10 h presented as bar charts with error bars representing standard deviation over 3 repeats. The fluorescence generated in the *P.aeruginosa* cultures was significantly higher than that produced by any other bacterial species (p = 0.006).

In this study, 7-hydroxycoumarin (7HC) was selected as the fluorescent reporter because of its favourable physicochemical and photophysical properties, including high chemical stability, straightforward synthesis, excellent photostability, high fluorescence quantum yield, and relative pH independence ^18,19^.The aminopeptidase catalyses hydrolysis of the peptide bond, resulting in cleavage of the amino acid recognition group and formation of an intermediate species (Steps 1 and 2). This intermediate then rapidly undergoes self-immolative 1,6-elimination of the PABA linker, leading to the release of the free fluorophore, 7-hydroxycoumarin (Step 3). Release of the fluorophore generates a fluorescence “turn-on” response, with emission observed at approximately 460 nm. The resulting increase in fluorescence intensity provides a direct and sensitive measure of aminopeptidase activity.

## Results and discussion

### Synthesis of fluorescent probes

Details of the synthetic route and design rationale used to generate the fluorescent probes is described in the supplemental data file. The method aims to establish a general protocol that can be used for multiple amino acids, dipeptides or other short peptides, but is validated here using a beta-alanine amino acid derivative.

### Microbiology assay development

The probe activation was measured using a continuous growth curve protocol, routed in the standard method for microbroth dilution, as described in the Material and Method section. This approach also allows the assay to be readily integrated with other microbiological methods, including minimum inhibitory concentration (MIC) assays and reduces assay variation. An input of approximately 5x10^5^ cfu/ml diluted from an overnight culture in cation-adjusted Mueller Hinton Broth was used throughout. The probe concentration was carefully optimised to ensure that activation could be monitored throughout a typical 20-hour bacterial growth curve without saturating signal, with bacterial strains that exhibited the highest levels of probe activation. This concentration was subsequently applied across all experiments, regardless of the bacterial species examined or the amino acid substrate incorporated into the individual probe design. The use of a standardised probe concentration enabled direct qualitative comparison of fluorescence signals throughout the study.

The probes were originally tested against a panel of strains used for the evaluation of novel antimicrobials as part of UKHSA’s Open Innovation in AMR platform ^20^. The strains represent a mix of relatively drug susceptible and drug-resistant strains, with known antibiotic resistance gene profiles and phenotypic susceptibility data, as shown in Table 1.

**Table 1:** ESKAPE panel strains used for initial probe activations studies. Abbreviations for common antibiotics to which strain are resistant AMX Amoxicillin AMP Ampicillin PIP Piperacillin GEN Gentamicin TOB Tobramycin CTX Cefotaxime CAZ Ceftazidime AMK Amikacin KAN Kanamycin CIP Ciprofloxacin NOR Norfloxacin MOX Moxifloxacin LVX Levofloxacin IMP Imipenem MEM Meropenem CST Colistin VAN Vancomycin

| Species | Strain name in thesis | Source | Drug Resistance Profile | Multidrug resistant (Resistant $\geq 3$ frontline classes of antibiotic) |
| --- | --- | --- | --- | --- |
| *K. pneumoniae* | KP13368 | NCTC13368 | AMX, AMP, PIP, GEN, TOB, CTX, CAZ | Resistant |
| *K. pneumoniae* | M6 | UKHSA | AMX, AMP, PIP | Sensitive |
| *A. baumannii* | AYE | ATCC BAA-1710 | AMX, AMP, GEN, AMK, KAN, TOB, CIP, NOR, CAZ | Resistant |
| *P. aeruginosa* | Ab17978 | ATCC 17978 | AMX, AMP | Sensitive |
| *P. aeruginosa* | PAO1 | NCTC | AMX, AMP, PIP | Sensitive |
| *P. aeruginosa* | PA13437 | NCTC13437 | AMK, AMX, CIP, NOR,<br>MOX, LVX, GEN,<br>AMK, KAN, TOB, CAZ,<br>IMP, MEM | Resistant |
| *E. coli* | EC12923 | NCTC12923 | AMX, GEN, TOB, AMK | Sensitive |
| *E. coli* | LEC001 | PHE | AMX, AMP, CIP, NOR,<br>LVX, CTX, CAZ, CST | Resistant |
| *S. aureus* | MSSA9144 | ATCC 9144 | – | Sensitive |
| *S. aureus* | MRSA-15 | NCTC13142 | CIP, NOR, AMX<br>(methicillin) | Resistant |
| *S. aureus* | MRSA-16 | NCTC13143 | CIP, NOR, AMX | Resistant |
| *E. faecalis* | NCTC 775 | NCTC775 | – | Sensitive |
| *E. faecalis* | NCTC12201 | NCTC12201 | VAN | Resistant |
| *E. faecium* | NCTC12204 | NCTC12204 | VAN, AMX |  |

Beta-alanine aminopeptidase activity has been reported to be selective for *P.aeruginosa* and has since been utilised in commercial chromogenic agar, i.e. ChromID® to test for P.aeruginosa infections in cystic fibrosis patients ^21^. In this study, the beta-alanine probe was activated strongly by both Pseudomonas strains while none of the other bacterial strains showed any significant fluorescence signal.

Beta-alanine 7HC probe was screened against 10 diverse strains of *P.aeruginosa* listed in Table 2, and compared with PAO1 as the comparator strain (Figure 3).

**Figure 3.**
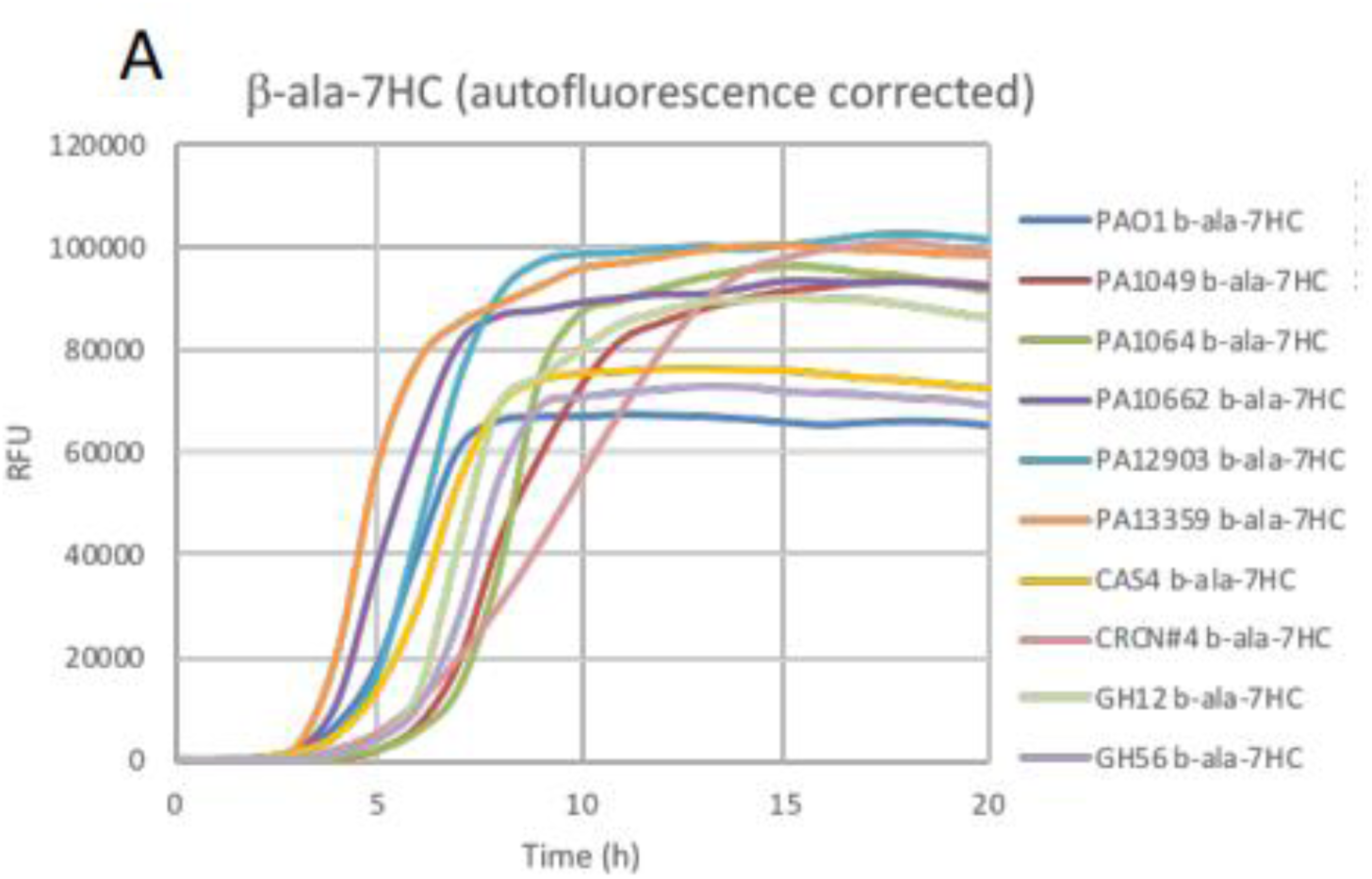
Fluorescence activity of the beta-alanine aminopeptidase probe after culture with P.aeruginosa strains shown in Table 1. The fluorescence assay is performed by adding diluted overnight bacteria culture of OD600 0.005 with 25µM of each probe on a microplate. The fluorescence generated was measured at 460 nm (7HC) at 37°C, hourly over 20 hours. OD600 curves (not shown) show that the growth of bacteria is similar in the presence and absence of probes, indicating no inhibitory effect of probes. The data was background corrected to remove the effects of autofluorescence of the bacterial strains, on an individual strain basis. Each graph is representative of at least 3 biological repeats with 2 technical replicates in each repeat.

**Figure 4.**
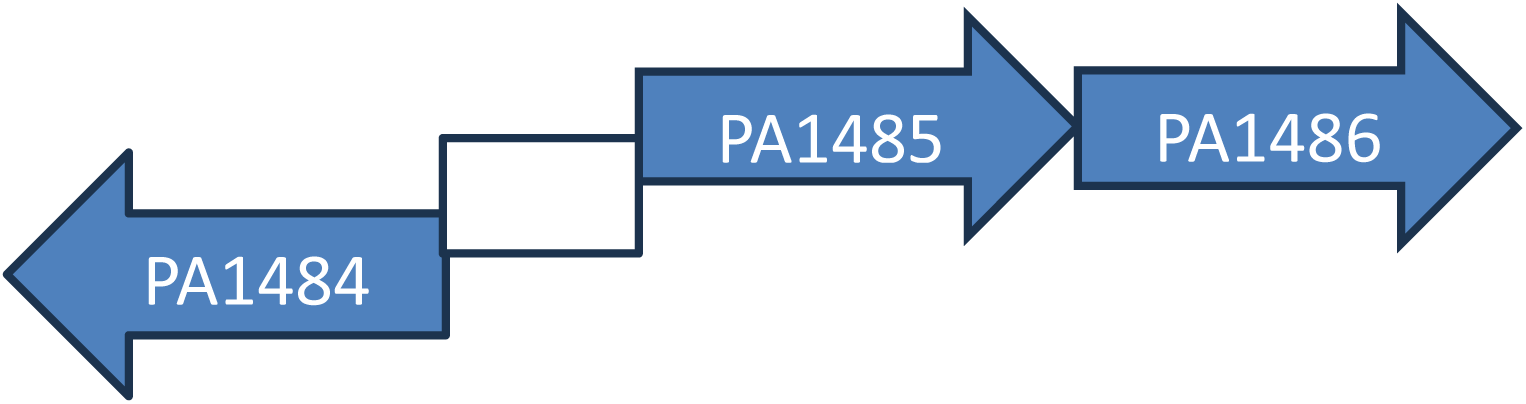
Diagrammatic representation of the *bapF* operon in PAO1 genome. The operon contains 3-genes, with a divergently expressed transcriptional regulator (PA1484), separated by a small intergenic region, from a co-expressed predicted permease (PA1485) and the bapF gene (PA1486). Operon structure is based on annotation of Gene ID: 881720 <u>PA1486 hypothetical protein [Pseudomonas aeruginosa PAO1] - Gene - NCBI</u>; updated on 30^th^ July 2025 and accessed on the 26^th^ June 2026. ^24, 25^

**Table 2.** Diverse panel of *P.aeruginosa* strains used in additional probe activation studies.

| Species/ strain | Strain label | Note; Source |
| --- | --- | --- |
| *P. aeruginosa* PAO1 | PAO1 | National Collection of Type Culture (NCTC) collection |
| *P. aeruginosa* 1049 | PA1049 | Avirulent; Outbreak strain from Birmingham hospital |
| *P. aeruginosa* 1064 | PA1064 | Avirulent; Outbreak strain from Birmingham hospital |
| *P. aeruginosa* NCTC 10662 | PA10662 | Drug-sensitive control strain; NCTC collection |
| *P. aeruginosa* NCTC 12903 | PA12903 | Drug-sensitive control strain; NCTC collection |
| *P. aeruginosa* NCTC 13359 | PA13359 | NCTC collection |
| *P. aeruginosa* CAS4 | CAS4 | Meropenem-resistant; Salisbury District Hospital |
| *P. aeruginosa* CRCN#4 | CRCN#4 | Carbapenem-resistant carbapenemase-negative strain; AMRHAI Colindale |
| *P. aeruginosa* GH12 | GH12 | Cystic fibrosis sample; UKHSA |
| *P. aeruginosa* GH56 | GH56 | Cystic fibrosis sample; UKHSA |

**Table 3.** List of transposon mutant strains used in this study, selected from PAO1 transposon mutant library.

| Strain Name | PA ORF | Putative ORF Function | Position in ORF |
| --- | --- | --- | --- |
| PAO1 | Wild type | – | – |
| PW3674 | PA1484 | probable transcriptional regulator | 394(807) |
| PW3676 | PA1485 | probable amino acid permease | 566(1368) |
| PW3678 | PA1486 | BapF | 630(1101) |

All tested strains of *P.aeruginosa* showed similar activation levels with the beta-alanine 7HC probe (Figure 3). There were some strain specific differences, with PA13359 and PA12903 showing more rapid activation to peak levels and a higher overall peak fluorescence than the control PAO1 strain. The time taken to show activation above the background fluorescence level was between 3 to 5 hours at the cell numbers a used, but there were again minor strain differences with PA1064 showing the slowest activation, compared to PA13359 which was quickest. The observed differences could relate to changes in the overall growth rate and hence cell numbers and/or the endogenous levels of enzyme expression and activity in individual strains.

To confirm that the observed activation was attributable to the annotated beta alanine aminopeptidase gene, variable described as *bapF* and *dmpA* , isolates from a transposon library of *P.aeruginosa* PAO1 were used.^22,23^

Results show that fluorescence probe activation was essentially eliminated in the *bapF* transposon mutant, (PW3678; PA1486; Fig 5A). This confirms that BapF is both essential and sufficient for beta-alanine aminopeptidase activity in *P.aeruginosa.* The data was further interpreted using integration to calculate the area under the curve (AUC) between T = 0 – 10h (Figure 5B). Since integration takes into account the time of activation, the gradient of the curve and the overall magnitude of the fluorescent signal, it better represents the total fluorescence activity. This confirmed that the probe activation was lost with the bapF transposon mutant. The integration data also showed that the transposon mutants in the putative transcriptional regulator and the predicted amino acid permease (PA1484 and PA1485, respectively) had a higher probe activation, reflecting the more rapid activation of the probe in the growth curve experiments. T-test shows that the fluorescence difference between control strain PAO1 with PW3674:PA1484, PW3676:PA1485 is statistically significant (P < 0.01). Time to half-maximal fluorescence also supported the observation of higher beta alanine aminopeptidase activity with values across the three repeat experiments averaging 6.74 hours for PAO1, but only 5.18 and 4.87 hours for PA1484 and PA1485 mutants respectively.

**Figure 5.**
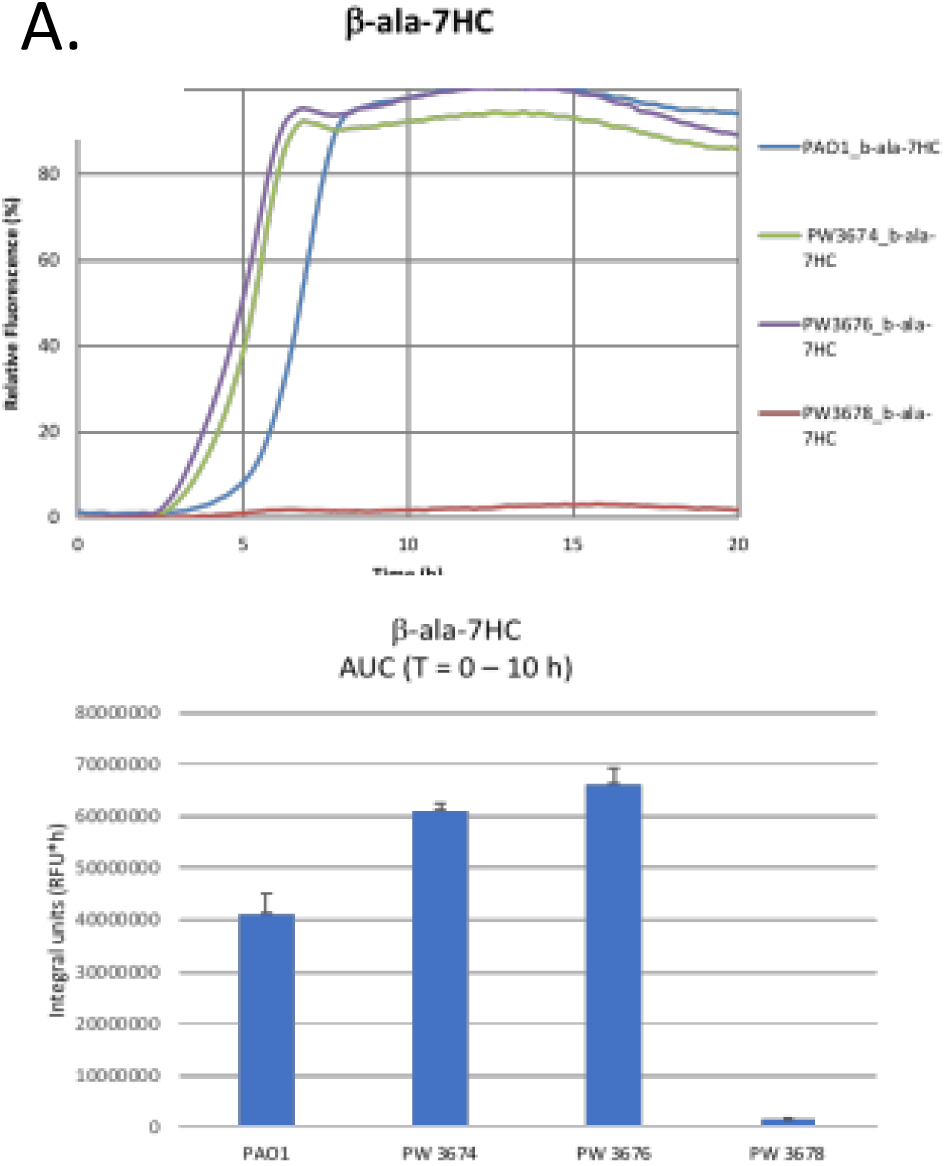
Fluorescence activity of b-ala-7HC when tested against PAO1 transposon mutants compared to the wild type PAO1. A. Normalise fluorescence data was generated from growth curves of PAO1 and three transposon mutants in PW3674, PW3676 and PW3678, respectively (PA1484, PA1485 and PA1486). Data is a representative of 3 repeats. B. Relative fluorescence was compared at T = 10 h. with data showing the average of 3 repeats and error bars representing the standard deviation.

A possible explanation for the observed phenotype is that the putative transcriptional regulator acts as a repressor of *bapF* expression, such that the transposon mutant de-represses expression, leading to higher production of the enzyme and more rapid probe activation. This might be linked to the phenotype of the permease, if there is a metabolic regulation of the repressor through binding of beta-alanine or beta-alanine containing peptides. In the absence of the putative permease, less beta-alanine gets into the cell so again there is de-repression of the transcriptional regulator, with elevated BapF production. This also assumes that the observed beta-alanine aminopeptidase probe activation activity is at least partly outside of the bacterial cell. There are other potential explanations, such as polar effects of the transposon insertions on *bapF* expression, which were not explored here. The potential role of the operon in beta-alanine scavenging and uptake is intriguing, perhaps providing *P.aeruginosa* with an alternative route to acquisition of beta-alanine as an essential cofactor for Coenzyme A ^26^, compared to the PanD-dependent synthetic route seen in other bacteria^27^.

In our initial testing of ESKAPE pathogen isolates, we only saw activation of the beta alanine probe in *P.aeruginosa*, but a number of other bacterial species are also b-peptidyl aminopeptidase producers (BAP). These include Sphingosinicella spp.^28,29^ *dmpA* from Ochrobactrum anthropic ^30^, BapA from Gram-positive bacterium Mycolicibacterium smegmatis, and BapA from two Burkholderia spp ^31^. All known representatives of this enzyme family are named b-peptidyl aminopeptidases (EC 3.4.11.24). Genome database shows that there are *bapF / dmpA*-homologous genes present in other bacterial species, although they have been not been fully characterised. We selected additional strains with predicted *bapF* genes (Table 4) for assessment with the beta-alanine aminopeptidase probe. This included 4 strains of Serratia marcescens, 7 strains of *Burkholderia cepacia* complex, and 4 strains of *Ochrobactrum anthropii* and 2 of *O.intermedium*. Fluorescence activation was measured essentially as described previously (Fig 6).

**Figure 6.**
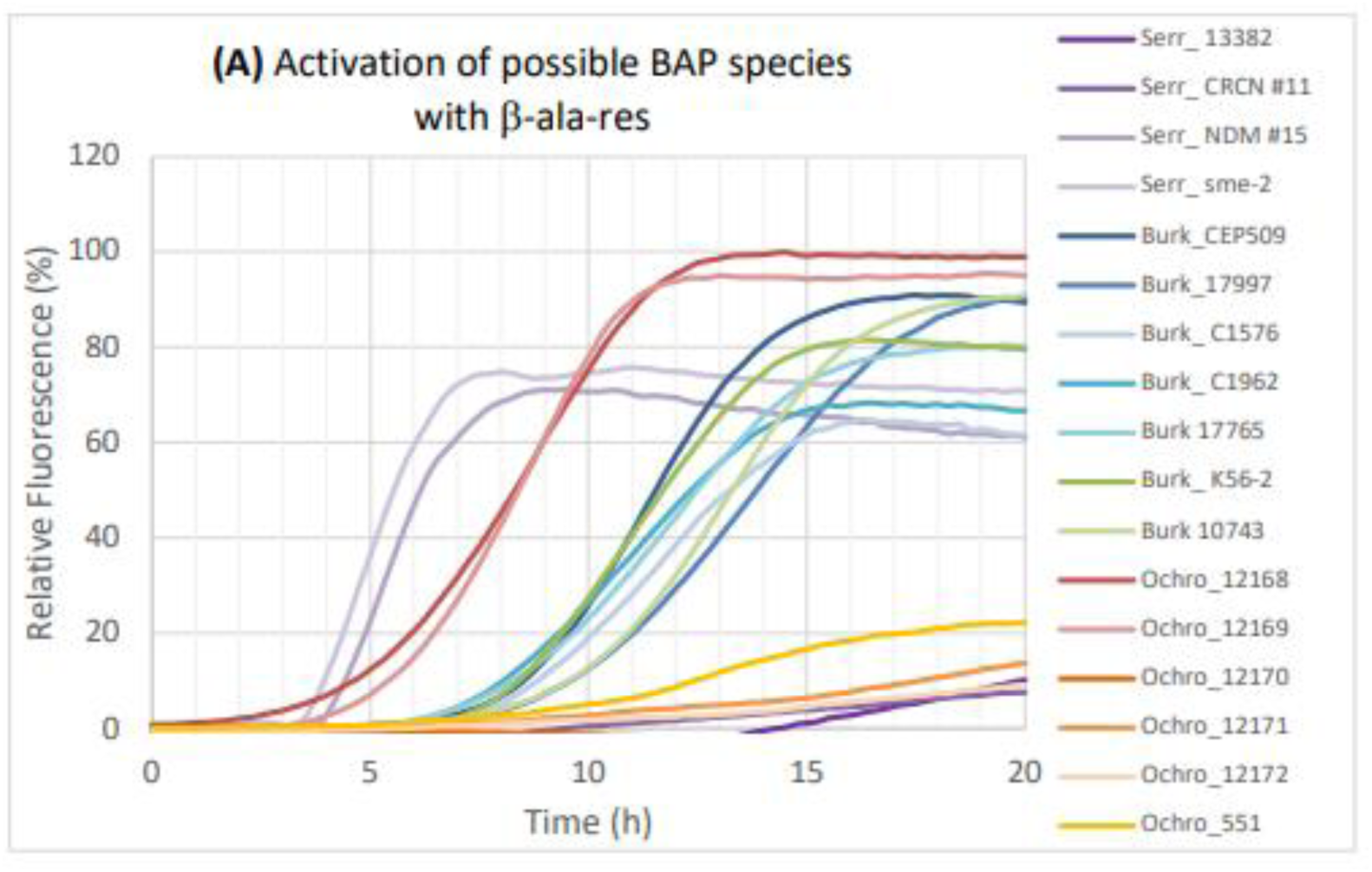
Beta-alanine aminopeptidase probe activation by different bacterial pathogens. Normalised fluorescence activity was observed in different strains of bacteria, measured as a 20-hour fluorescence growth curve, starting with approximately 5x10^5^ cfu/ml in Mueller Hinton Broth. Graphs are representative of 3 biological repeats with 2 technical replicates in each experiment.

**Table 4.** List of additional bacteria strains used for assessment of potential beta alanine aminopeptidase activity.vn.

| **Serratia spp.** | **Strain** | **Burkholderia spp.** | **Strain** | **Ochrobactrum spp.** | **Strain** |
| --- | --- | --- | --- | --- | --- |
| *S. marcescens* | 13382 | *B. cepacia* | 509 | *O. anthropi* | NCTC12168 |
| *S. marcescens* | CRCN#11 | *B. cepacia* | 17997 | *O. anthropi* | NCTC12169 |
| *S. marcescens* | NDM#15 | *B. cepacia* | 17765 | *O. anthropi* | NCTC12170 |
| *S. marcescens* | sme-2 | *B. cepacia* | 10743 | *O. anthropi* | 551 |
|  |  | *B. cenocepacia* | K56-2 | *O. intermedium* | NCTC12171 |
|  |  | *B. multivorans* | C1576 | *O. intermedium* | NCTC12172 |
|  |  | *B. multivorans* | C1962 |  |  |

For *S.marcescens*, only two strains NDM#15 and sme-2 are able to activate the beta alanine aminopeptidase probe, with both strains showed activation between 3 to 4 hours, with fluorescence increasing rapidly to reach maximal fluorescence around 8 hours. Strain CRCN#11 and 13382 showed no activation until 10 – 14 hours where there was only a limited amount of probe activation. This may reflect late expression of the beta alanine aminopeptidase or that release of the enzyme from the cell is only observed in stationary phase.

All seven strains of *Burkholderia cepacia* complex (four strains of *B.cepacia*, one strain of *B.cenocepacia,* and two strains of *B.multivorans*) showed activation of the probe with activation seen around 6-7 hours, reflecting slower overall growth of these strains.

Activation was strain dependent with two of the four strains of *O.anthropi* (12168, 12169) showing very rapid probe activation, with activation observed within 1 hour, whilst the other two strains showed limited activation (strain 551) or no activation (12170). No activation was observed for the two strains of *O.intermedium* (12171, 12172). All of the Ochrabactrum spp. isolates had genes annotated as *dmpA* (data not shown), but sequence differences between the genes suggest that some of these may be misannotated, have different peptide substrate specificities and/or have very different levels of expression under the conditions tested.

## Conclusions

We show here a general scheme that can be applied to the generation of switch on aminopeptidase probes, exemplified by beta alanine, but with the potential to be used for natural or non-natural amino acids, dipeptides or other short peptides. The probe was applied to confirm both species and enzyme selective nature of activation, by beta alanine aminopeptidases in *P.aeruginosa* and a limited range of other species tested.

By examining the predicted operon structure in PAO1 and leveraging transposon mutants to evaluate changes in phenotype, we were able to confirm that the annotated *bapF* gene was responsible for probe activation and identified a possible beta-alanine responsive transcriptional regulator and a identified a possible beta-alanine responsive transcriptional regulator and an amino acid permease that may transport beta alanine and/or beta-alanine containing peptides. The potential of this operon to play a role in scavenging beta alanine in different physiological settings is intriguing, especially given its conservation in Burkholderia species, both species being associated chronic lung infections. Further studies will investigate the significance of this conserved system as well as exploiting the beta-alanine 7HC probe’s ability to monitor growth of *P.aeruginosa* in essentially real time, in complex microbiome or tissue extract.

## Materials and Methods

### Synthesis of beta-alanine 7HC probe

Design principles and beta-alanine aminopeptidase probe synthesis is described in detail in the supplementary information file.

From the commercially available Boc-β-ala-OH and 4-aminobenzyl alcohol, general procedure for amide coupling using EDCI and HOBt was used. The crude product was purified using column chromatography with 20 – 50 % acetone in DCM (v/v), affording pure product (1.120 g, 46.9% yield) as a pale yellow solid.

**1 H NMR** (400 MHz, CDCl3+MeOD) δ ppm 8.92 (s, 1H), 7.46 (d, J=8.44Hz, 2H), 7.25 (d, J=8.44Hz, 2H), 5.50 (s, 1H), 4.57 (s, 2H), 3.37 (t, J=5.92Hz, 2H), 2.48-2.52 (m, 2H), 1.40 (s, 9 H);

**13C NMR** (101 MHz, CDCl3+MeOD) δ ppm 170.24, 156.70, 137.31, 136.84, 127.69, 120.07, 119.97, 79.80, 64.56, 49.60, 49.39, 49.18, 37.28, 28.33.

### Biological Evaluation

#### General Materials and Methods

All chemicals and reagents used were of molecular grade and obtained from Sigma-Aldrich unless otherwise stated. Bacterial strains used in this thesis are described in Chapter 6.2.2. All strains were maintained on Tryptic Soy Agar (TSA) plates (BioMérieux) and overnight cultures are grown in cation-adjusted Muller-Hinton broth II (MH2) or Tryptic Soya broth (TSB) unless otherwise stated. Overnight cultures were prepared by selecting a single colony of bacteria from a TSA plate using a sterile loop and added into a universal flask filled with MH2 media (3 mL). For Enterococcus spp., TSB media was used instead of MH2 for the overnight cultures due to poor overnight growth in MH2.

Bacterial culture was incubated at 37 °C for 12 – 18 h in a shaking incubator at 200 rpm. The optical density (OD) of the overnight culture was determined using a spectrophotometer the next day and diluted to the desired OD to be used in experiments. Unless otherwise stated, the starting OD used in all screening experiments is OD600= 0.01. The growth of bacteria was monitored by measuring the optical density at 600 nm (OD600) on the plate reader. The synthesised fluorescent probes are stored in glass vials or in an Eppendorf vial in the freezer at -20 °C. A stock solution of 20 mM for each probe was prepared by dissolving known mass of the solid compound with calculated volume of DMSO and are stored in the freezer. Before using in experiments, the compounds are diluted from the stock solution to the desired concentration in MH2 media.

#### Real time fluorescence growth curves

A single colony was picked from a TSA plate and grown in 3 mL of media overnight in a shaking incubator at 37 °C. The following day, the OD600 of the overnight culture was measured and diluted to OD600 = 0.01 in media. On a 96-well plate, in each well of final volume 200 *μ*L, bacteria of starting OD600 = 0.01 was added 100 *μ*L per well, followed by the addition of 100 *μ*L per well of the compound with starting concentration of 50 *μ*M, to give 25*μ*M final probe concentration, unless otherwise stated. For control wells, where no probe or no bacteria was needed, 100 *μ*L of MH2 broth was added instead. For every plate, there are three types of control well: media-only control well, bacteria-only well, and probe-only well. The 96- well plate was covered with a lid and incubated at 37 °C on the plate reader (FLUOSTAR Omega BMG Labtech) for 20 h during which absorbance and fluorescence were continually measured every hour. Absorbance was measured at 600 nm, and fluorescence was measured at two wavelengths, *λ ex*/*em* = 355/ 460 nm for 7HC compounds, and *λ ex*/*em* = 540/590 nm for resorufin compounds. The plate reader’s gain setting was set at optimised setting for the experiments.

#### Data analysis

All fluorescence data are either representative of three independent repeats (without error bars) or the average of three biological repeats (with standard deviation as error bars). Using Microsoft Excel or MARS data analysis software (BMG Labtech) the average of two technical replicates in each biological repeat was calculated and plotted in Excel to give a 20 h fluorescence line graphs. Data from control wells were plotted to ensure that there is no contamination, and the fluorescence from the experimental wells were blanked against the background fluorescence from the probe-only control wells. The integration or area under the curve values were calculated using MARS data analysis software, and the values are tabulated using Excel spreadsheet to calculate the average and standard deviation of three biological repeats. The data is plotted as bar charts and the standard deviation is represented by the error bars. P-values are calculated using two sample T-test function in Excel. P < 0.05 is statistically significant, and P < 0.005 is highly statistically significant. For growth curves, the optical density at 600 nm (OD600 ) values were plotted using Excel to produce growth curves of each bacteria strain. The growth curves of bacteria-only control are compared to the absorbance of experimental wells to investigate if the probes have any inhibitory effect on the growth of bacteria.

## Acknowledgements

We acknowledge technical support from Sirine Zadi, Carrie Turner and Ivy Zhu. Transposon mutants were obtained from the University of Washington^23^, the Manoil Group at Washington University Genome Services through funding from the NIH (P30 DK089507).

