## supplementary data for "Evaluation of selectively-activatable, caged fluorescent probes as species selective markers for beta-alanine aminopeptidase positive bacterial species"

**General Chemistry methods.**

Unless otherwise stated, all glassware were dried overnight on a drying rack or oven dried for 15 – 30 min. Reactions carried out at ambient room temperature is at 20 – 30°C. The temperature of 0°C was achieved using an ice bath, - 45 °C using dry ice with acetonitrile, - 78 °C using dry ice with acetone. Reaction under inert atmosphere is by means of positive pressure of nitrogen or argon gas with a filled balloon or using the Schlenk line. Reaction progress were monitored using TLC- Thin Layer Chromatography performed on precoated Merck TLC silica gel 60 F254 aluminium sheets and visualized using UV light (short and long wave), and when necessary stained with potassium permanganate or vanillin prepared according to standard protocols. Purification of compounds using flash column chromatography was carried out using silica gel (Merck 9385, 230-400 mesh ASTM, 40-63 µM) as the stationary phase, or using a Grace Reveleris® X2 automated Flash Chromatography System. Removal of solvents or concentrated in vacuo is by the means of house vacuum connected to Buchi rotary evaporator, or KNF RC-600 evaporating system equipped with a SC-920 G vacuum pump (KNF). All solvents including anhydrous solvents are reagent grade solvents provided by Sigma-Aldrich, Acros, Alfa Aesar and Fisher Scientific, and are used as provided. Yields refer to isolated material (homogeneous by TLC or NMR) unless otherwise stated. Compound names are assigned according to IUPAC nomenclature. 1 H NMR was recorded on NMR Bruker AMX400 (400MHz). 13C NMR was recorded on NMR Bruker AMX400 (101MHz). NMR data were reported in the order: chemical shift (ppm), multiplicity, coupling constant (Hz), integration and assignment of peaks when possible. Abbreviations for multiplicities are as follows: s=singlet, d=doublet, t=triplet, q=quartet, m= multiplet, dt= double triplet, td= triple doublet. 2D NMR such as COSY, DEPT, HMBC spectra were also run to further confirm the structure of compound when necessary. High Resolution Mass Spectra (Exactive HCD Orbitrap mass spectrometer, Thermo Scientific) was acquired using electron ionisation or chemical ionisation technique depending on the substrate. All Liquid Chromatography Mass Spectroscopy (LCMS) analysis was performed on a Waters Alliance 2695 with water (A) and acetonitrile (B) comprising the mobile phases. Formic acid (0.1%) was added to both acetonitrile and water to ensure acidic conditions throughout the analysis. Function type: Diode array (535 scans). Column type: Monolithic C18 50 X 4.60 mm. Mass spectrometry data were collected using a Waters Micromass ZQ instrument coupled to a Waters 2695 HPLC with a Waters 2996 PDA. Waters Micromass ZQ parameters used were: Capillary (kV), 3.38; Cone (V), 35; Extractor (V), 3.0; Source temperature (°C), 100; De-solvation Temperature (°C), 200; Cone flow rate (L/h), 50; Desolvation flow rate (L/h), 250. LCMS gradient conditions are described as follows. Method A (10 min): from 95 % A/ 5 % B to 50% B over 3 min. Then from 50 % B to 80 % B over 2 min. Then from 80% B to 95% B over 1.5 min and held constant for 1.5 min. This was then reduced to 5% B over 0.2 min and maintained to 5% B for 1.8 min. The flow rate was 0.5 mL/min, 200 µL was split via a zero dead volume T piece which passed into the mass spectrometer. The wavelength range of the UV detector was 220-400 nm. Method B (5 min): from 95 % A/ 5 % B to 90% B over 3 min. Then from 90% B to 95% B over 0.5 min and held constant for 1 min. This was then reduced to 5% B over 0.5 min. The flow rate was 1.0 mL/min, 100 µL was split via a zero dead volume T piece which passed into the mass spectrometer. The wavelength range of the UV detector was 220-500 nm.

***Amide coupling using HATU and DIPEA***

Boc-protected amino acid (1.0 equiv.) in DCM (5 mL) was added with DIPEA (2.0 equiv.) and HATU (1.2 equiv.). The suspension was stirred at r.t. for 15 min under N2 gas before the addition of para-aminobenzyl alcohol, PABA (1.1 equiv.). The reaction was stirred at r.t.

overnight and the progress of reaction was monitored using TLC and/or LC-MS. Upon completion, the reaction was diluted with DCM (15 mL) and washed with saturated $NaHCO_3$ (3 x 15 mL), 1M citric acid (3 x 15 mL), deionised water (2 x 15 mL), and finally brine (2 x 15 mL) to remove aqueous-soluble impurities. The organic layer was dried over anhydrous $MgSO_4$, filtered and concentrated in vacuo to yield the crude product which was then further purified using either flash column chromatography, recrystallisation or hot filtration method as stated.

***Mesylation (Chlorination)***

Using anhydrous conditions, starting material (1.0 equiv.) in dry DCM at 0 oC was added with dry $Et_3N$ (2 equiv.) and then mesyl chloride (2 equiv.) under inert atmosphere. Reaction was stirred for 2 h under inert atmosphere at r.t. Reaction progress was monitored using TLC, and another portion of dry $Et_3N$ (2 equiv.) and mesyl chloride (2 equiv.) were added again under ice bath. Upon completion, the reaction was worked up by diluting with DCM and the washing of organic layer with deionised water (3 x 15 mL), saturated $NaHCO_3$ (3 x 15 mL), and brine (3 x 15 mL). The organic layer was dried over $MgSO_4$, filtered and concentrated in vacuo to afford the crude product which was either used as it is, or purified using flash column chromatography.

**Substitution reaction with 7HC fluorophore**

Starting material (1.0 equiv.) was added with 7-hydroxycoumarin, 7HC (1.0 equiv.) and $K_2CO_3$ (2.0 equiv.) in DMF at r.t. Reaction was stirred at r.t. overnight. DMF was removed in vacuo before working up using saturated $NaHCO_3$ (3 x 15 mL), deionised water (3 x 15 mL) and brine (3 x 15 mL). The organic layer was dried over $MgSO_4$, filtered and concentrated in vacuo to afford the crude product which was either used as it is, or purified using flash column chromatography. Substitution reaction with resorufin fluorophore In a small round-bottomed flask or microwave vial wrapped with aluminium foil, resorufin sodium salt (0.9 equiv.) and K2CO3 (2.0 equiv.) in DMF was stirred at r.t. before adding starting material (1.0 equiv.). Reaction was stirred at r.t. overnight. DMF is removed in vacuo after LCMS showed completion of reaction. EtOAc was added and the organic layer was washed with copious amount of deionised water and brine until the aqueous layer ran clear. The organic layer was dried over anhydrous MgSO4, filtered and concentrated in vacuo to give product, which was used directly in the next step, or purified using reverse phase column chromatography due to its polar nature.

***Deprotection of Boc group using TFA***

Starting material (1.0 equiv.) was added with a solution of 1:1 TFA-DCM (v/v) and stirred at r.t. for 20 mins at 0 oC. Reaction was monitored using TLC. Upon completion of reaction, equivalent volume of toluene was added before concentrating in vacuo to yield the final product as a TFA salt.

***Deprotection of Boc group using HCl***

Starting material (1.0 equiv.) was dissolved in minimal MeOH (< 2 mL) and then added with 4 M-HCl in dioxane (1 mL) at 0 oC. Reaction was stirred at r.t. for 20 min at r.t. and was monitored using TLC. Upon completion of reaction, equivalent volume of toluene was added before concentrating in vacuo to yield final product as a HCl salt.

***Synthetic approach and methods:***

The designed retrosynthetic approach II was reversed and translated into a synthetic route to the target molecule (Scheme 2.5.1) which forms the basis of our synthesis. By utilising a different starting block, 4-aminobenzyl alcohol, this new synthetic route eliminated the ester reduction step described in the literature. With a reduction in number of synthetic steps, fluorescent probes can now be synthesized in a shorter time, and the use of hazardous reducing agents can be avoided altogether.

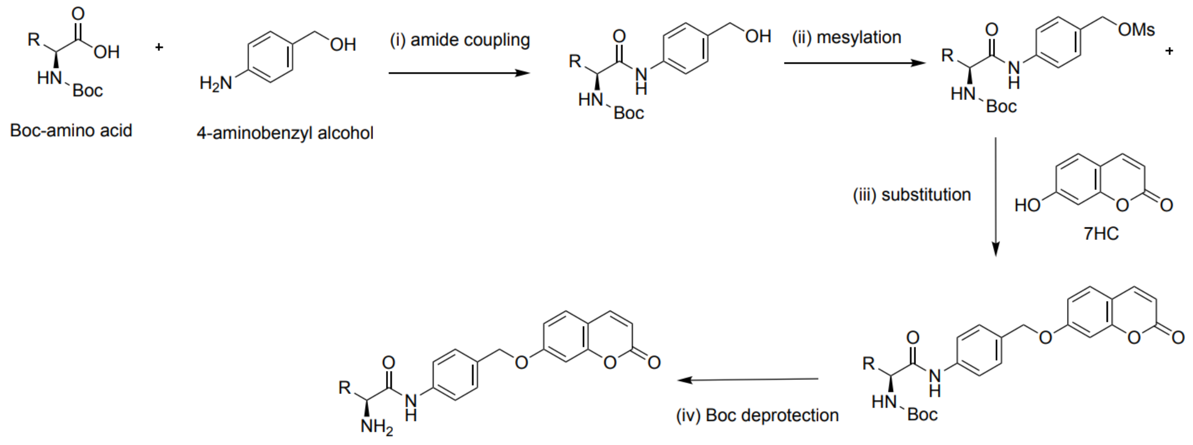


Figure S1; General synthetic route. R= amino acid side chain

This step-economical synthesis also means that the associated use of other reagents and solvents used in reaction, work-up and purification, together with its waste could be reduced. Scheme 2.5.1 General synthetic route. R= amino acid side chain. O HN N H R O OMs HN N H R OH O HN H2N R OH OH Boc Boc-amino acid 4-aminobenzyl alcohol HO O O 7HC O HN N H O O O O R NH2 N H O O O R (i) amide coupling (ii) mesylation (iii) substitution (iv) Boc deprotection Boc Boc Boc 49 2.5.2 Amide coupling reaction The general synthetic route begins with the formation of an amide bond between N-Bocprotected amino acid and commercially available 4-aminobenzyl alcohol. Scheme 2.5.2 Amide coupling reaction to form intermediates 2.2- 2.10 (b) The amide coupling step was initially carried out using amide coupling reagents EDCI and HOBt as with the synthesis of the reference compound L-ala-7HC. During an amide coupling reaction, the carboxylic acid is activated with stoichiometric amounts of EDCI which converts the OH into a better leaving group, forming the intermediate O-acylisourea. The intermediate then reacts with HOBt to form a HOBt ester and corresponding urea as a by-product. The second step is the acylation of the amine group by the HOBt ester to give the corresponding amide product, and regeneration of HOBt. The use of HOBt as an additive is to optimise reactivity and to prevent racemisation. HOBt also prevents the formation of undesired Nacylurea, which can occur as a result of rearrangement from O-acylisourea. In an attempt to increase reaction yields, and minimise laboratory risk since HOBt is potentially explosive, HATU was used instead for the synthesis of later compounds. Although the improvement in yields were not major, the reaction was more convenient with HATU. The addition of HOBt into the reaction vessel needed to be carried out at 0 °C using an ice-water bath as a precautionary measure, whereas the HATU reaction can be carried out at room temperature. Moreover, when EDCI-HOBt was used, an additional step was required to filter off-white precipitate formed during the work-up. The mechanism of HATU is similar to EDCIHOBt as HATU reacts with carboxylic acid to form OAt-active esters which then reacts with amine to afford the product. O HN N H R OH O HN H2N R OH OH Boc Boc-amino acid 4-aminobenzyl alcohol (i) amide coupling Boc 2.2-2.10 (b) 50 2.5.3 Mesylation (Chlorination) reaction Since alcohols are poor substrates for substitution reactions as the hydroxyl group is a strong base, hence poor leaving group, the hydroxyl group needs to be converted to a better leaving group in order to facilitate the subsequent SN2 reaction. Sulfonates such as methanesulfonate (Mesylate, OMs) and p-toluenesulfonate (Tosylate, OTs) are very good leaving group. Scheme 2.5.3 Mesylation reaction of compounds 2.2- 2.10 (b) to give the intermediates 2.2-2.10 (c) This reaction is achieved using methanesulfonyl chloride (MsCl), and triethylamine in anhydrous conditions. Triethylamine catalyses the reaction by acting as a base, deprotonating the intermediate hence speeding up the reaction. It also removes HCl as it forms during the reaction, to make sure excess HCl that forms do not affect the acid- labile Boc protection group. After an aqueous work up, compounds were either used without further purification, or purified via column chromatography on silica gel. The leaving group intermediates were checked using LC-MS followed by NMR. Interestingly, the spectral data for the intermediate compounds do not corroborate with the formation of mesylate. The LC-MS results did not show the expected mass of the mesylated compounds. Instances where 1 H NMR were performed to confirm this observation, the NMR spectra lacked the methyl peak of the mesylate group, which is usually at around 3.0 to 3.5 ppm, or even anywhere else on the spectra. Upon further investigation, LC-MS analysis showed that for all compounds, the chlorinated compound was formed instead. The molecular weight of the chlorinated compounds and their LC-MS peaks are showed in Table 2.5.1. The LC-MS data also showed the characteristic 3: 1 ratio of the molecular ion peaks M+ and M+2, which reflects the fact that chlorine isotope 35Cl is three times more than the isotope 37Cl. Thus, it is evident that the chlorination reaction had occurred instead of a mesylation reaction. This observation is further supported by reported literature that during the tosylation of alcohol using tosyl O HN N H R O OMs HN N H R OH (ii) mesylation Boc Boc 2.2-2.10 (b) 2.2-2.10 (c) 51 chloride, the chlorinated product instead of tosylated product was formed. 157 A plausible explanation offered by the authors is that the tosylate intermediate is further displaced by the nucleophilic chloride ion to afford the chlorinated counterpart (Scheme 2.5.4). Scheme 2.5.4 Plausible mechanism of the formation of chlorinated product 2.2-2.10 (c) Since halogens are also good leaving groups, and the chlorinated intermediate worked well in the following substitution reaction, thus no changes to the synthetic route were made. Table 2.5.1 LC-MS analysis of the intermediates (b) of Aminopeptidase library I, supporting the formation of chlorinated product but not the mesylated product.

***Purification and characterisation of compounds***

In most cases, the purification of the intermediates was achieved using column chromatography on silica gel to afford compounds with adequate degree of purity for characterisation and for use in subsequent reactions or microbiological evaluation. A mixed solvent system of EtOAC/hexane or EtOAc/DCM with varying degree of polarity was found to be the most efficient in separating impurities from the product for these reactions. For amide coupling reactions, an acidic work-up using 1M citric acid solution was performed to remove any unreacted amines. In cases where the purity levels are fairly low after workingup, recrystallisation or filtration using hot solvent 1:1 EtOAc- Hex (v/v) was carried out. If there is more than one spot on the TLC or LCMS, then column chromatography was carried out instead. The intermediates and final compounds have been fully characterised and reported in the chemistry experimental section in Chapter 6. Intermediates are characterised using LC-MS (5 min method) and 1 H NMR, with the occasional use of other methods such as LC-MS (10 min method), 13C NMR and/or HR-MS when necessary. The final compounds are characterised using LC-MS (5- and 10-min method), 1 H NMR, 13C NMR, and HR-MS methods. Overall, the synthesis of probes had been achieved with moderate- good yields to give final compounds in high purities suitable for use in biological evaluation. Methods including NMR, LCMS and HRMS have been used to fully characterised the intermediates and final compounds (Chapter 6). 55
